# Drowsiness during resting-state fMRI: a major confounder of functional connectivity

**DOI:** 10.1101/2023.06.19.545521

**Authors:** Marc Joliot, Sandrine Cremona, Christophe Tzourio, Olivier Etard

## Abstract

This research explores the effects of drowsiness on variability in functional connectivity (FC) during resting-state functional magnetic resonance imaging. The study utilized a cohort of students (MRi-Share) and classified individuals into drowsy (N=68), alert (N=96), and mixed/undetermined states based on observed respiratory oscillations. Five different processing methods were employed, the reference method, two correction methods based on physiological and global regression approaches, and two based on Gaussian standardizations. According to the reference methodology, the results indicate that drowsy individuals exhibit higher cortico-cortical FC than alert individuals. However, the differences between drowsy and alert states were reduced when applying correction methods based on physiological and global regression approaches. The global regression-based strategy was the most effective among these correction methods, minimizing significant FC differences to only 3.3% of the total FCs. Utilizing the Gaussian-based methods, both cortico-subcortical and intra-default mode network regions demonstrated significantly greater FCs in awake than drowsy subjects. These findings align with previous studies suggesting that, in the descent to sleep, the cortex isolates itself to facilitate the transition into deeper sleep stages while also disconnecting the default mode network. The Gaussian standardization methods and the global regression-based correction approach efficiently address the hemodynamic variations caused by the rapid alternation between the N1 stage and wakefulness. These variations contribute to the measurement of cortico-cortical pseudo connectivity observed in the reference methodology. In summary, these findings underscore the importance of considering drowsiness in rs-fMRI studies and demonstrate that there is no single optimal correction methodology for processing fMRI data

## Introduction

Functional imaging during resting state, also known as the brain default mode (Andreasen et al., 1995, Mazoyer et al., 2001, Raichle et al., 2001), refers to a behavioral and physiological state experienced when deprived of meaningful cognitive stimulation. Utilizing functional magnetic resonance imaging (fMRI), researchers have observed regional hemodynamic low-frequency pseudo-oscillations (LFOs) (Biswal et al., 1995), which exhibit similarity between distant brain regions. Subsequent investigations have revealed that this signal extends throughout the cortical, subcortical, and cerebellar gray matter. Moreover, the functional connectivity (FC, see review van den Heuvel and Hulshoff Pol, 2010)), denoting the signal similarity between regions, has emerged as an indicator of the cognitive anatomo-functional organization of the brain [Smith, 2009].

The resting state encompasses two types of processes. The first involves homeostatic mechanisms likely associated with synaptic activity maintenance (Laumann and Snyder, 2021). The second process is also linked to synaptic activity but is related to ongoing cognition during fMRI acquisition. Notably, these two processes are not mutually exclusive, as demonstrated in earlier studies by Grecius et al. (Greicius et al., 2003). The pseudo-periodic nature of LFOs suggests a stronger association with the first process. Indeed, investigations have explored and observed modulations of functional connectivity in relation to genetic/developmental factors (e.g., sex), learning/cognitive skills, and neuropsychiatric brain diseases [(See reviews of Vaidya and Gordon, 2013 and Canario et al., 2021 for the latter). The second process is self-oriented activity and a mental state associated with mind-wandering (Antrobus et al.,). In imaging studies, it has been described as “random episodic silent thinking (REST)” (Andreasen et al., 1995), “default mode” (Raichle et al., 2001), or “conscious resting state” (Binder et al., 1999, Mazoyer et al., 2001). Capturing this “random” process is challenging since it involves a continuous flow of thoughts without any external triggers. Nevertheless, certain studies have reported variability in functional connectivity associated with the content of these thoughts (Andrews-Hanna et al., 2010; Doucet et al., 2012; McKeown et al., 2020; Smallwood et al., 2016; Wang et al., 2018). However, even when investigating life events or ongoing cognition, the linked variability in functional connectivity accounts for only a tiny portion of the observed variance (Doucet et al., 2012). It is important to note that fMRI acquisitions are subject to various sources of variability, and determining which factors should be accounted for remains an ongoing topic of research and debate. In addition to the sources mentioned above of functional connectivity variability, four undesired sources exist: instrumental artifacts, subject movement-related artifacts, non-specific tissue artifacts, and physiological artifacts. Each source requires specific correction methodologies, comprehensively reviewed by Caballero-Gaudes et al. (Caballero-Gaudes and Reynolds, 2017).

During the analysis of 15-minute resting state (RS) functional magnetic resonance imaging (fMRI) data from the MRi-Share cohort (Tsuchida et al., 2021), a database consisting of 1722 students and acquired using a 3 Tesla MRI scanner, we observed substantial remaining variability in FC even after applying the correction mentioned above methods. They were instructed to keep their eyes closed to increase the likelihood of subjects entering a mind-wandering state. However, as Tagliazucchi et al. (Tagliazucchi and Laufs, 2014) demonstrate, such a state promotes drowsiness. Hence, it raises the question of whether FC variability could be influenced by the subject’s entry into or avoidance of this drowsy state. In addition to diminished vigilance, drowsiness is likely to induce physiological changes in the brain’s signals, potentially impacting the intrinsic connectivity observed during the resting state.

Numerous changes have been observed when studying the transition from wakefulness to sleep using rs-fMRI. The global blood oxygen level-dependent (BOLD) signal, measured across the entire brain, tends to increase as sleep deepens (McAvoy et al., 2019), whereas global cerebral blood flow measured by other techniques, such as positron emission tomography, decreases concurrently (Braun et al., 1997). These observations collectively suggest a neurovascular decoupling induced by sleep. Although brain networks revealed by rs-fMRI analysis are present across all sleep stages (Koike et al., 2011), the strength of connectivity within the networks decreases during the deepest stage, and specific sub-networks are no longer observed (Horovitz et al., 2009). However, this decrease is not a continuous phenomenon along the wake-sleep axis, as studies have shown an increase in global connectivity during the initial phase of sleep (Larson-Prior et al., 2009; Tagliazucchi et al., 2012). Furthermore, the temporal dynamics of brain networks are altered by changes in alertness, with transitions between functional connectivity states occurring more frequently during wakefulness compared to the deepest sleep stages (El-Baba et al., 2019; Stevner et al., 2019). The lightest sleep stage is distinct as it does not correspond to any single state or group of states identified through dynamic analysis (Stevner et al., 2019).

In summary, studies investigating sleep using MRI techniques are limited due to their inherent complexity, but they consistently reveal changes in brain connectivity compared to wakefulness. These changes do not form a continuous spectrum across the wake-sleep axis, leading us to hypothesize that they can be observed as soon as the initial signs of drowsiness appear, a state that frequently occurs during an rs-fMRI scan (Tagliazucchi and Laufs, 2014).

The primary objective of this study was to distinguish, within our extensive dataset, individuals who experienced drowsiness from those who remained alert during the fMRI acquisition. To achieve this, we utilized the understanding that during sleep, not only do the electrical activities of neurons change but also autonomic regulation (Duyn et al., 2020). Apart from a decrease in the frequency of heartbeats and respiration, falling asleep gives rise to respiratory disturbances, including reduced chemoreceptor sensitivity and delayed feedback loops. These disturbances result in respiratory control system instability and periodic breathing characterized by alternating short episodes of hyperventilation and hypoventilation (Khoo et al., 1982). Although the exact occurrence of these phenomena as individuals fall asleep is unknown, we leveraged the opportunity presented by our large dataset, all of which was accompanied by photoplethysmography (Allen, 2007) and respiratory belt measurements, to categorize subjects into three groups: those experiencing drowsiness, those remaining fully alert, and those with a mixed or undetermined state.

Our analysis proceeded in two main steps. First, we examined the differences FC associated with drowsiness versus the alert state. Subsequently, we tested two methods of time series corrections and two methods of FC corrections for the observed differences. The initial two correction methods were based on the Physiological Response Function (PRF) (Kassinopoulos and Mitsis, 2019), including RETROICOR (Glover et al., 2000) and Global Normalization (GR), which involved regression using the entire BOLD time course of the grey matter. The other two methods directly corrected individual FCs by applying z-scores based on two distribution approaches.

## Materials and Methods

### Data acquisition

The MRI datasets used in this study were obtained from a subset of the i-Share (internet-based Student Health Research, www.i-share.fr) protocol, which aims to investigate the health of university students. Of the 30,000 enrolled students in i-Share, 1870 volunteered to participate in the MRI study, forming the MRi-Share cohort (Tsuchida et al., 2021). The MRi-Share study was conducted with the approval of the local ethics committee (CPP2015-A00850-49). After excluding subjects who did not complete the scanning session and those with incidental findings, a final sample of 1722 subjects remained for further analysis.

From the three acquired modalities, namely anatomical, diffusion, and functional MRI, we extracted ten sequences for this study. The T1-weighted images were obtained using a sagittal 3D magnetization-prepared 180-degree radio-frequency pulses and rapid gradient-echo (MP RAGE) sequence with the following parameters: repetition time (TR)/echo time (TE)/inversion time (TI) = 2000/2.0/880 msec and a voxel size of 1mm^3^. For fMRI geometric distortion correction, we utilized eight B0 diffusion mapping sequences as described in (Tsuchida et al., 2021). The resting-state fMRI acquisition consisted of a 15-minute scan with 1058 volumes using a 2D axial Echo Planar Imaging (EPI) sequence (TR/TE = 850/35.0 msec, Flip Angle = 56°, voxel size 2.43 mm^3^). This acquisition was performed on a 3T Siemens scanner with a 64-channel head coil, utilizing the “Minnesota” multi-band sequence with six times acceleration. Among 1640 subjects, who underwent the resting-state acquisition, simultaneous recordings of two physiological signals were acquired: blood volume changes in the microvascular bed of finger tissue using photoplethysmography (PPG, Allen, 2007) and breathing movements using a pneumatic abdominal respiration transducer. During the resting state scan, participants were instructed to keep their eyes closed, relax, avoid movement, remain awake, and allow their thoughts to come and go freely. Compliance with these instructions was verified using a self-report questionnaire administered at the end of the resting state scanning session (Cremona et al., 2022).

### Physiological Signal Processing and Vigilance Assessment

The physiological signals of the 1640 subjects underwent initial visual screening to identify acquisition artifacts. We identified two main types of artifacts: periods of signal loss and clipping that affected either one or both signals. Overall, this resulted in the exclusion of 270 subjects from further analysis. The second visual analysis focused on the respiratory signal to identify a characteristic pattern of drowsiness, characterized by alternating waxing and waning in the amplitude of abdominal respiratory movements. Previous research by Trinder et al. demonstrated that these amplitude alternations are associated with transitions between wakefulness and N1 sleep stage, with decreases in ventilation during wakefulness to N1 state transitions and increases during N1 to wake state transitions (Trinder et al., 1992). Therefore, for the group of subjects exhibiting oscillatory breathing indicative of drowsiness, their respiratory signal envelopes had to show clear modulation of more than 50% of the amplitude with a period of approximately 30 seconds (Khoo et al., 1982) over more than half of the resting test duration (as shown in the first curve of Figure 1). In contrast, for the group of alert subjects, the respiratory signals were required to exhibit minimal envelope modulation with a period of about 30 seconds (as shown in the second curve of Figure 1). Based on these criteria, we identified 164 participants out of the initial 1370 as either showing patterns of drowsiness (N=68) or alertness (N=96) based on their respiratory patterns (Figure 1).

**Fig. 1:**
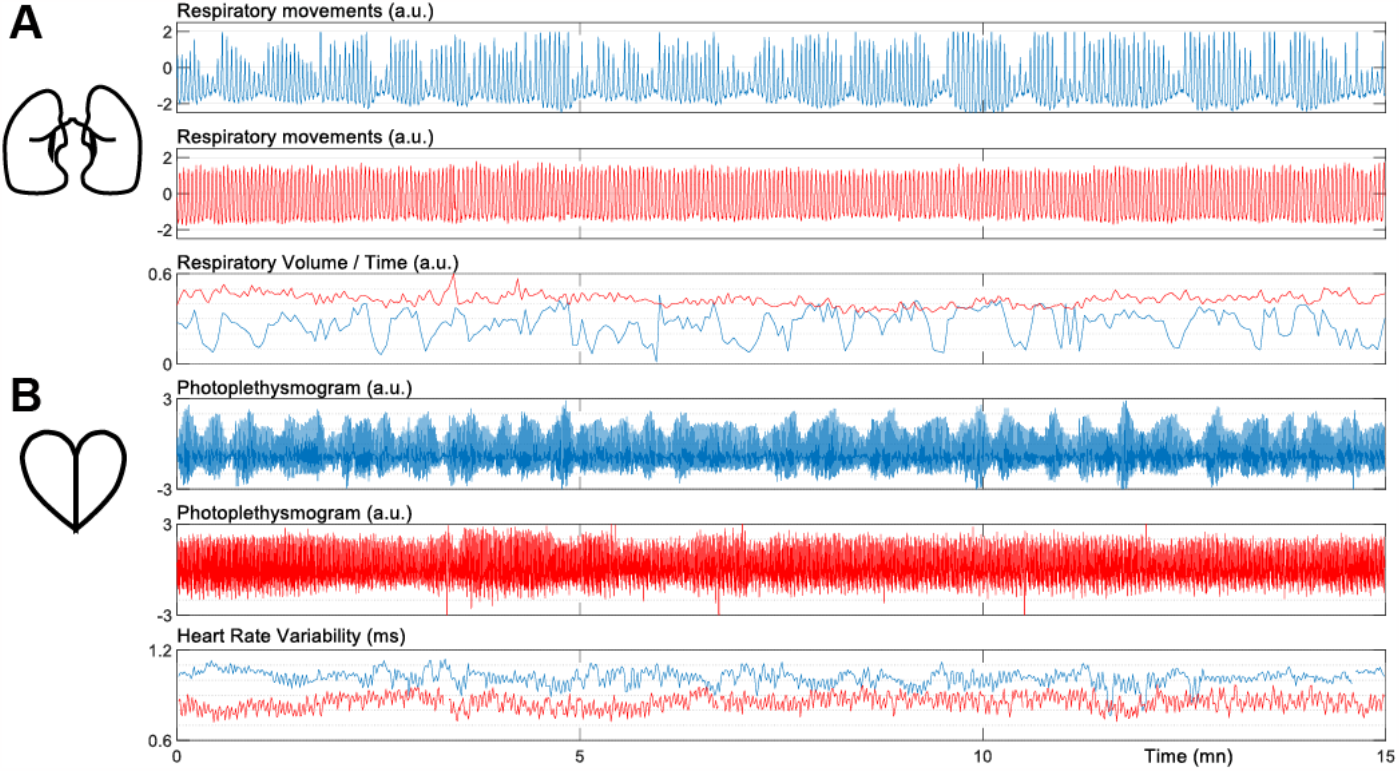
Respiratory (A) and heart (B) patterns of typical drowsy (blue) and alert (red) subjects.

We conducted a group analysis to characterize the physiological variables of these two populations. Initially, we automatically detected each breath and heartbeat using the PhysIO toolbox (Kasper et al., 2017). Subsequently, all detected events were individually checked and manually corrected, if necessary, using custom software. The following variables were extracted: the time difference between two consecutive breaths on the respiratory movements signal to estimate instantaneous breathing rate and the estimated Respiratory Volume per Time (RVT) (Figure 1.A bottom). Similarly, the time difference between two consecutive cardiac beats on the PPG signal was used to estimate instantaneous heart rate (i.e. the tachogram), and heart rate variability (Figure 1.B bottom) was assessed based on this tachogram. Finally, frequency analysis was performed on the RVT and tachogram using a multitaper spectrogram (Prerau et al., 2017).

### Resting-state processing

The preprocessing of rs-fMRI data followed state-of-the-art standards outlined in Tsuchida et al. 2021 (Tsuchida et al., 2021). First, using T1-weighted acquisition, normalization to the MNI template and segmentation into three tissue classes (GM: Gray Matter, WM: White Matter, CSF: Cerebrospinal Fluid) was performed. To ensure signal stabilization and eliminate subject responses to experimental onset, such as scanner noise, the initial 33 seconds of EPI data were removed. The remaining EPI volumes (1023) underwent spatial registration, and B0 acquisitions were utilized to compute a B0 field map, which was then employed to correct for geometrical distortions in the EPI images. The average registered and corrected EPI data were further registered to the T1-weighted image, and each EPI volume was interpolated into the MNI stereotactic space. The time courses of the six parameters describing movement across time and the average BOLD signal in the eroded WM and CSF were extracted. Two control quality markers, head motion (mHD: mean relative displacement, Alfaro-Almagro et al., 2018)) and mean Framewise Displacement (mFD: mean of the instantaneous FD, Esteban et al., 2019, Power et al., 2012), were derived from the movement parameters to quantify motion in two ways. A set of nuisance variables was constructed using the six movement parameters and their derivatives, the WM and CSF BOLD time courses, and sine and cosine time series corresponding to temporal filtering frequencies below 0.01 Hz. Subsequently, temporal filtering and nuisance regression were performed in a single regression model to minimize potential errors in data denoising (Caballero-Gaudes and Reynolds, 2017) using AFNI 3dREMLfit. This analysis was referred to as the Reference analysis (REF) after that. The FC between each pair of regions (384 regions based on the AICHA atlas, Joliot et al., 2015) was computed as the Pearson correlation coefficient between the average time series of the voxels belonging to the respective regions. This resulted in the computation of approximately 73,000 FCs in each of the 164 subjects. Finally, from the resting state questionnaire, we extracted the question regarding awareness of sleepiness (QAS), which was answered on a four-level scale (“never, rarely, often, and all the time”).

### Correction of Drowsiness Effect

Four methods were tested to account for differences in FCs between subjects experiencing periods of drowsiness and those fully alert. The first two methods were applied to the time courses, while the last two were applied directly to the FC values. The first method involved constructing regressors of non-interest based on the two acquired physiological signals, using both RETROICOR (Glover et al., 2000 and Physiological Response Function (PRF, Kassinopoulos and Mitsis, 2019). This method will be referred to as Ret-PRF. The PRFsc or M04 scan-specific model was used to estimate the cardiac and respiratory response functions. The correction level was evaluated by computing the correlation between the whole brain time courses and the estimated physiological response functions for both the heart and respiratory signals.

The second methodology employed global regression (GR) through regression using the whole brain BOLD time course. The third method involved z-score normalization of each subject’s FC distribution (Zs). In contrast, the fourth method implemented z-score normalization of the Gaussian of a mixture modeling of the FC distribution (Gaussian + two gamma functions), referred to as ZMix.

### Statistical analysis

To assess the significance of vigilance (drowsiness: N=68, alertness: N=96) for the FCs computed based on the reference and four correction methods, the quality control markers for movement (mHD and mFD), and the QAS extracted from the resting state questionnaire, ANOVA components were performed using a mixed-effect linear regression model. Random effects were fitted at the participant level, and the analyses were adjusted for age, sex, and brain volume. Significant differences were reported at a p-value threshold 0.05 for the quality control markers and QAS, and Bonferroni correction was applied for the FCs (N=384*383/2=73536). Additionally, the results were summarized for each AICHA region by calculating the percentage of significant effects (with 100% representing 383).

### Synthesis of the FC differences across the five methodologies

Firstly, we combined the results of the five analyses of variance (ANOVA). We constructed square FC matrices of size 384×384 for each methodology. These matrices represented the significance of the FC difference between two states: drowsy and alert. We assigned a value of 1 if the FC was higher in the drowsy state compared to the alert state, -1 if it was higher in the alert state compared to the drowsy state, and 0 otherwise. Secondly, the five FC matrices were then concatenated, and a hierarchical decomposition was computed using JMP software (ward cost function). We divided the data into four classes, corresponding to cortical partitions in the sensory, executive, and Default Mode regions and deep nucleus classes.

Lastly, we quantify the FC within and between each class, we defined a metric called FC classe (FCc). The FCc was calculated as the difference between the number of significant FCs where drowsy was greater than alert and the number of significant FCs where alert was greater than drowsy. Each FCc was normalized by the total number of FCs to scale its value between -1 and 1.

## Results

### Auto evaluation and quality marker

In the auto-evaluation, a higher frequency of reporting awareness of sleepiness was observed in the drowsy group compared to the alert group (t=-5.16, df = 162, p < 0.001). The two movement quality control markers used in the fMRI analysis did not show a significant difference in mHD (t=-1.80, df = 162, n.s.) between vigilance levels. Still, drowsiness significantly increased mFD (t=-1.99, df = 162, p = 0.048).

### Physiological Analysis

The results of the physiological variable analysis are summarized in the upper part of Figure 2. Regarding respiration, drowsy subjects exhibited a significantly lower respiratory frequency (t=3.58, df = 162, p < 0.001) and Respiratory Volume per Time (RVT) (t=9.53, df = 162, p < 0.001) than the alert subjects. Frequency analysis of the RVT signal revealed that the continuous component was more pronounced in the alert subjects. In contrast, the drowsy subjects showed more oscillations around 0.033 Hz, confirming the distinction between the two groups based on breathing patterns (oscillatory vs. stable breathing). Regarding heart rate, the drowsy subjects exhibited a generally lower heart rate (t=4.18, df = 153, p < 0.001) and significantly higher cardiac variability (t=-4.30, df = 153, p < 0.001) than the alert subjects, particularly around the frequency of respiratory oscillations.

**Fig. 2:**
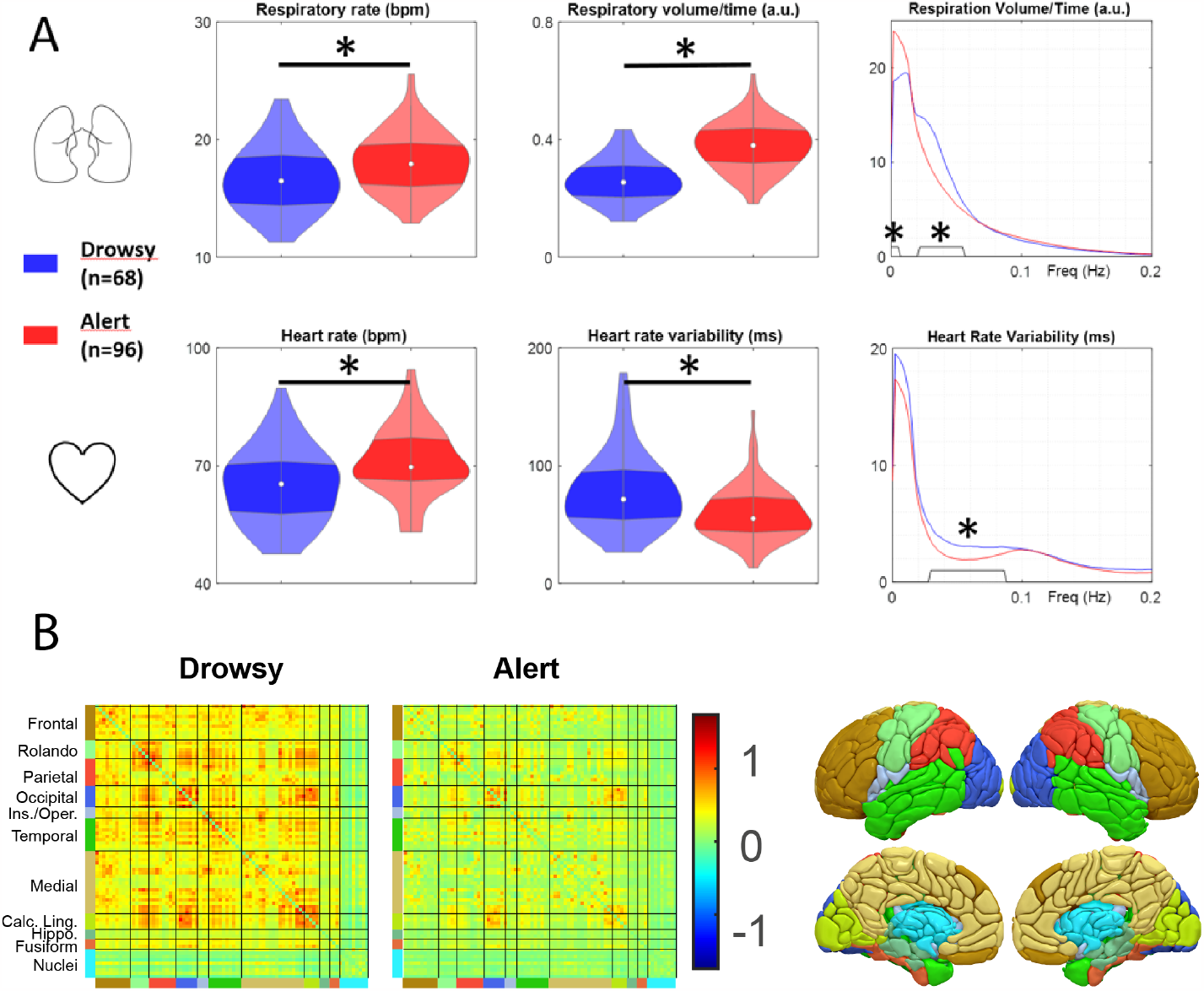
A) Analysis of respiratory rate, volume per time, and frequency content (top) and heart rate, rate variability, and frequency content (bottom) for drowsy and alert subject groups (*: p < 0.05). B) Group average intrinsic connectivity matrices computed in REF analysis. The color on the border of the matrices represents the areas shown in the 3-dimensional view of the atlas.

The correlation between the whole brain time course and the PRF Respiratory-based correction was found to be greater in the drowsy group (z-corr_Dr_ = 0.54 ± 0.13) compared to the alert group (z-corr_Al_ = 0.35 ± 0.13) (t_Dr-Al_ = 8.76, p < 0.0001, df=162). Conversely, the correlation for the PRF heart-based correction was higher in the alert group (z-corr_Al_ = 0.36 ± 0.14) compared to the drowsy group (z-corr_Dr_ = 0.28 ± 0.13) (t_Dr-Al_ = -2.83, p = 0.004, df=162).

### fMRI Reference Analysis

The group-average FCs are depicted in Figure 2.B. It reveals that cortical FCs are positive and higher in the drowsy group compared to the alert group. Moreover, FCs with most deep nuclei are lower than cortico-cortical FCs and similar in both groups.

Figure 3 illustrates the qualitative evolution of individual and group average FCs histograms. With the REF analysis, drowsy subjects shift their histograms towards high correlation values. The Ret-PRF analysis reduces this shift and the width of both average group distributions. The GR technique centers both distributions around null correlation values while maintaining a slightly more pronounced spread of positive values in the drowsiness group. The Zs technique increases the spread of the alert subject group towards positive values, and the ZMix technique further enhances this effect. Quantification of the group differences (t-ratio) based on the first four moments (mean, standard deviation, skewness, and kurtosis) of the individual histogram distributions are shown in Figure 3.F. The average absolute t-ratio of the four moments is minimal for the GR correction (see Figure 3.F dashed curve).

**Fig. 3:**
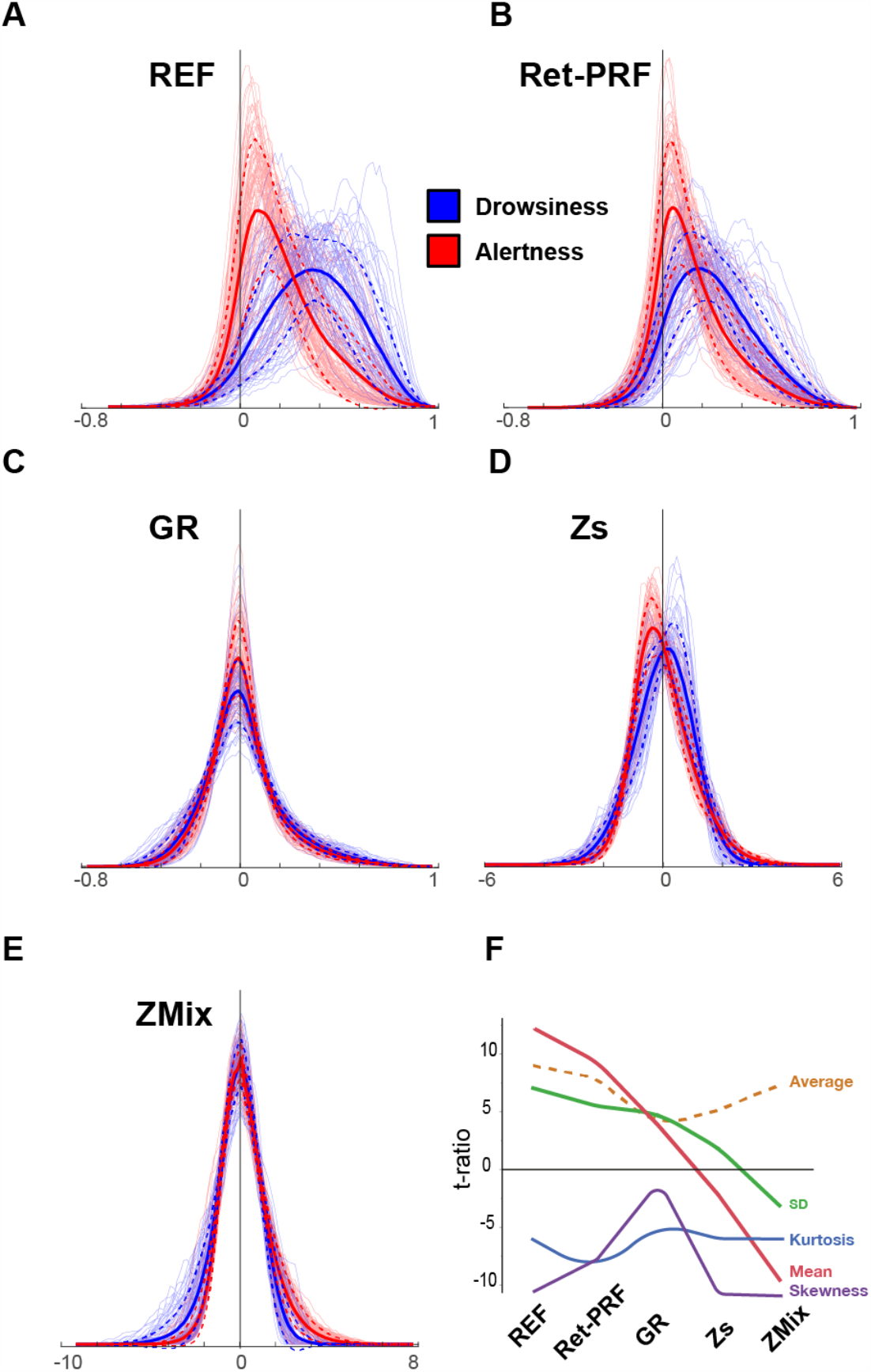
Histograms of inter-regional intrinsic connectivity for the five analyses: A) Reference (REF) analysis, B) Ret-PRF, C) GR (correction applied on the time course), and D) Zs and E) ZMix (correction applied on the FC values). The thick lines represent each class’s average (Drowsy, Alert) subjects, and the dashed line represents the average plus or minus one standard deviation. The thin lines represent individual subjects. F) T-ratio group differences of the first four moments of the individual histogram modeling. The dashed line represents the average of the absolute values of the four moments.

Figure 4 and Table 1 present the quantitative analysis of significant differences between drowsy and alert subjects in the REF analysis and the four correction methods. In the REF analysis, 65% of FCs were higher in drowsy subjects than alert subjects, while none showed the reverse pattern. This percentage decreased to 31% with the Ret-PRF correction and dropped below 5% for the other three techniques. Conversely, the number of alert above drowsy FCs remained zero with the Ret-PRF analysis but increased from 1% in GR to 6% in Sz and 18% in the ZMix analysis. The GR analysis had the lowest number of significant FCs at 3.3%.

**Table 1:** Percentage of the significant (Bonferroni corrected) vigilance effect on the inter-regional intrinsic connectivity.

| Type of Correction | Alert > Drowsy (%) | Drowsy > Alert (%) | Sign. FCs (%) |
| --- | --- | --- | --- |
| REF | 0 | 65.9 | 65.9 |
| Ret-PRF | 0 | 30.7 | 30.7 |
| GR | 1.0 | 2.3 | 3.3 |
| Zs | 6.2 | 4.4 | 10.6 |
| ZMix | 18.3 | 0.8 | 19.2 |

**Fig. 4:**
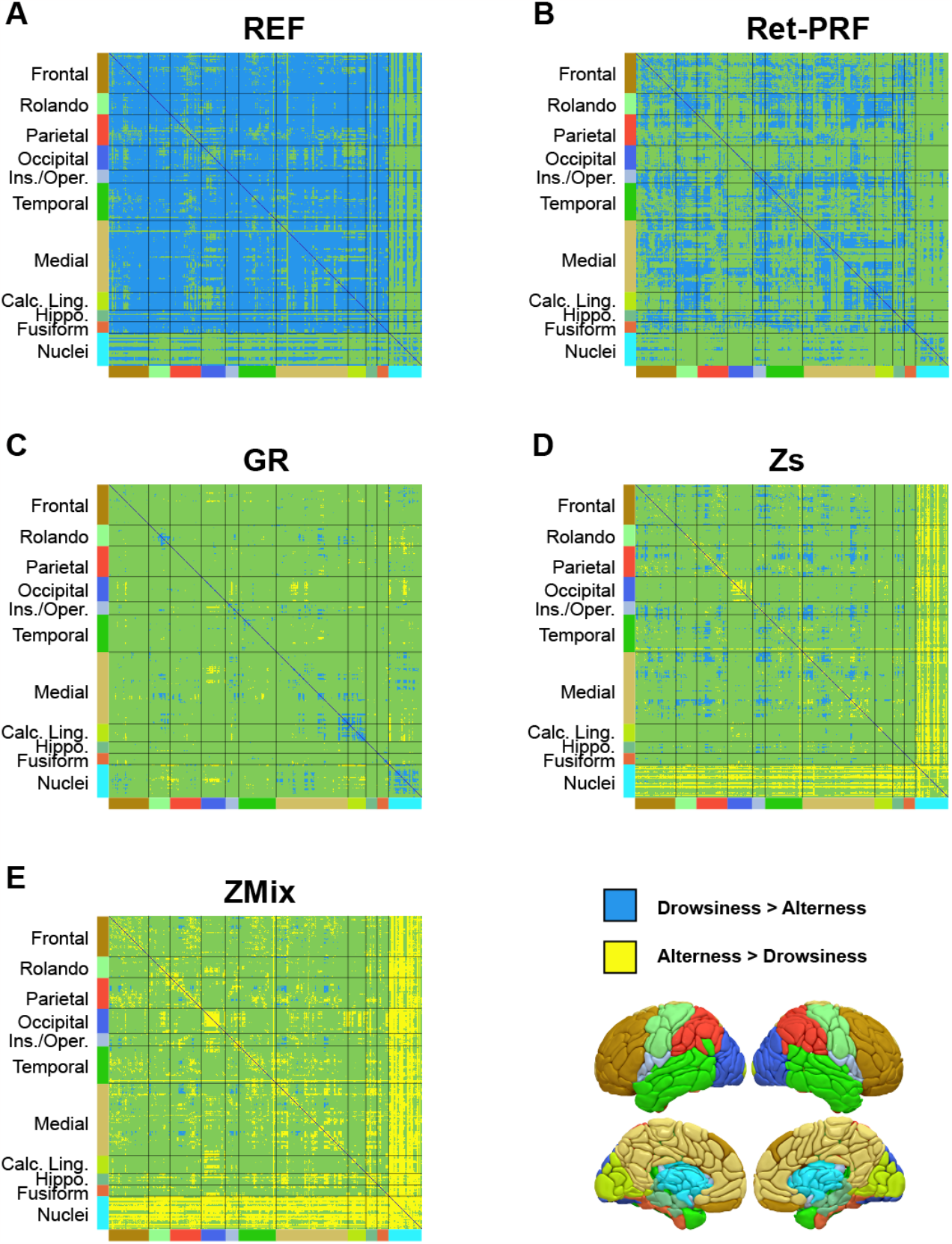
Significant effect of vigilance on inter-regional intrinsic connectivity calculated using the AICHA atlas (384 ROIs) for different analysis approaches: A) Reference analysis (REF), B) Ret-PRF correction, C) GR correction, D) Sz correction, and E) ZMix correction. The green represents ROI pairs with no significant difference, while blue and yellow indicate significant differences (p < 0.05 Bonferroni corrected). The color bordering the matrices corresponds to the areas depicted in the lower-left three-dimensional view of the atlas.

The regional specificity of these differences is illustrated in Figure 5 for the left hemisphere. The effect was observed without correction in all cortical regions, with higher magnitudes in the frontal areas and lower magnitudes in the occipital lobes. Both Figure 4A and Figure 5A demonstrate that most nuclei did not show any significant differences. The Ret-PRF-based correction reduced the vigilance effect in all cortical regions, while the GR-based correction further diminished it. Only with the GR correction, 0.7% of FCs were higher in alert subjects than in drowsy subjects, primarily involving FCs between deep nuclei and sensory-related cortical regions. These surviving FCs accounted for at least 30% of deep nuclei. With Sz and ZMix corrections, the percentage of FCs specific to the alert state increased, reaching 18%. Figure 4E (yellow bands) and Figure 5E (the hot-colored regions in the medial faces) illustrate that the FCs involving at least one deep nucleus captured the specificity of the alert state.

**Fig. 5:**
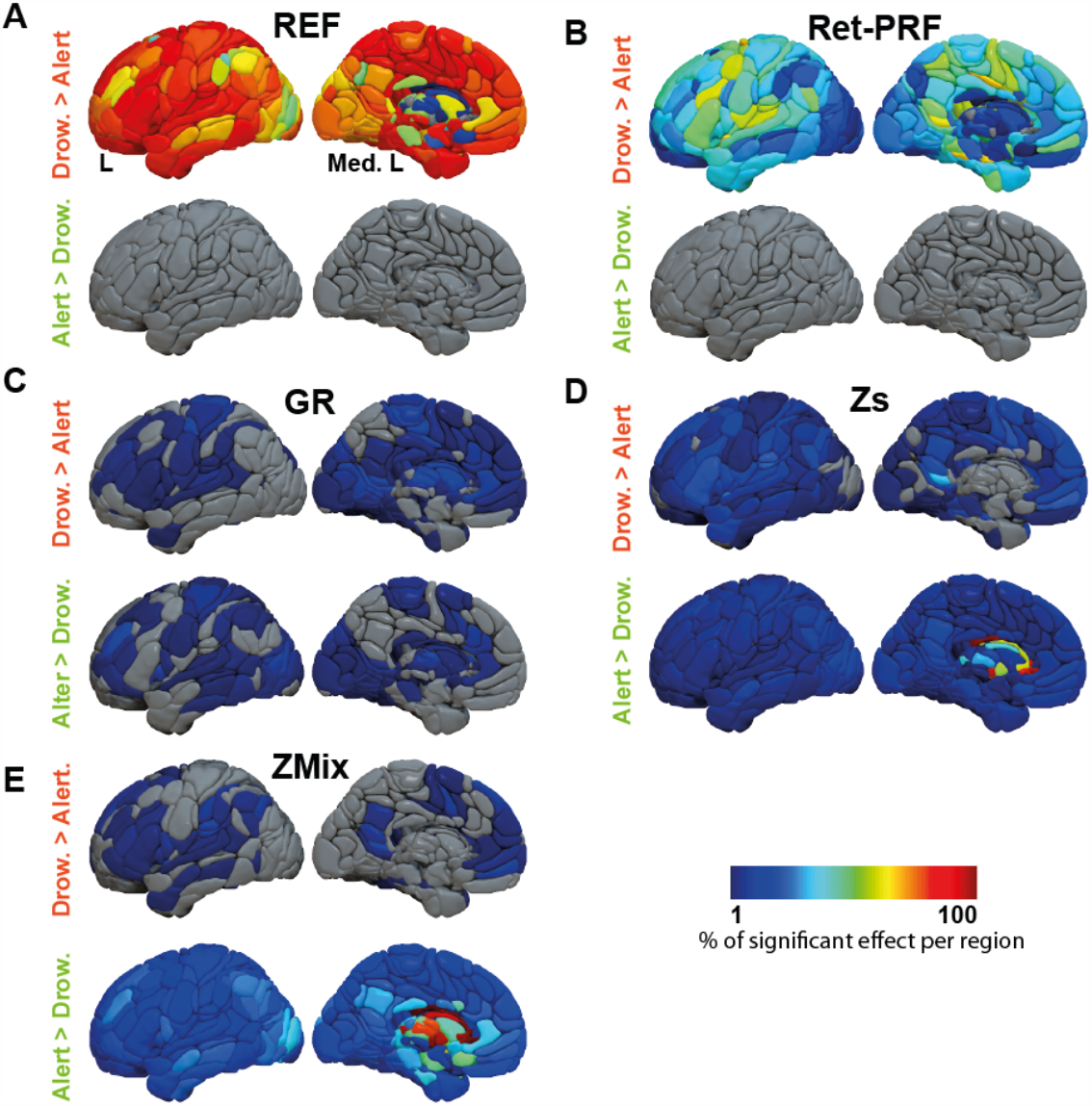
Synthesis of the vigilance effect on inter-regional intrinsic connectivity per region (only the left hemisphere is presented) and analysis approaches: A) Reference analysis (REF), B) Ret-PRF correction, C) GR correction, D) Sz correction, and E) ZMix correction.

Figure 6 displays the average FCs matrices for drowsy and alert subjects in the GR and ZMix corrections. As expected, the GR-derived maps looked similar, while the Mix analysis alert map exhibited larger values than the drowsy maps.

**Fig. 6:**
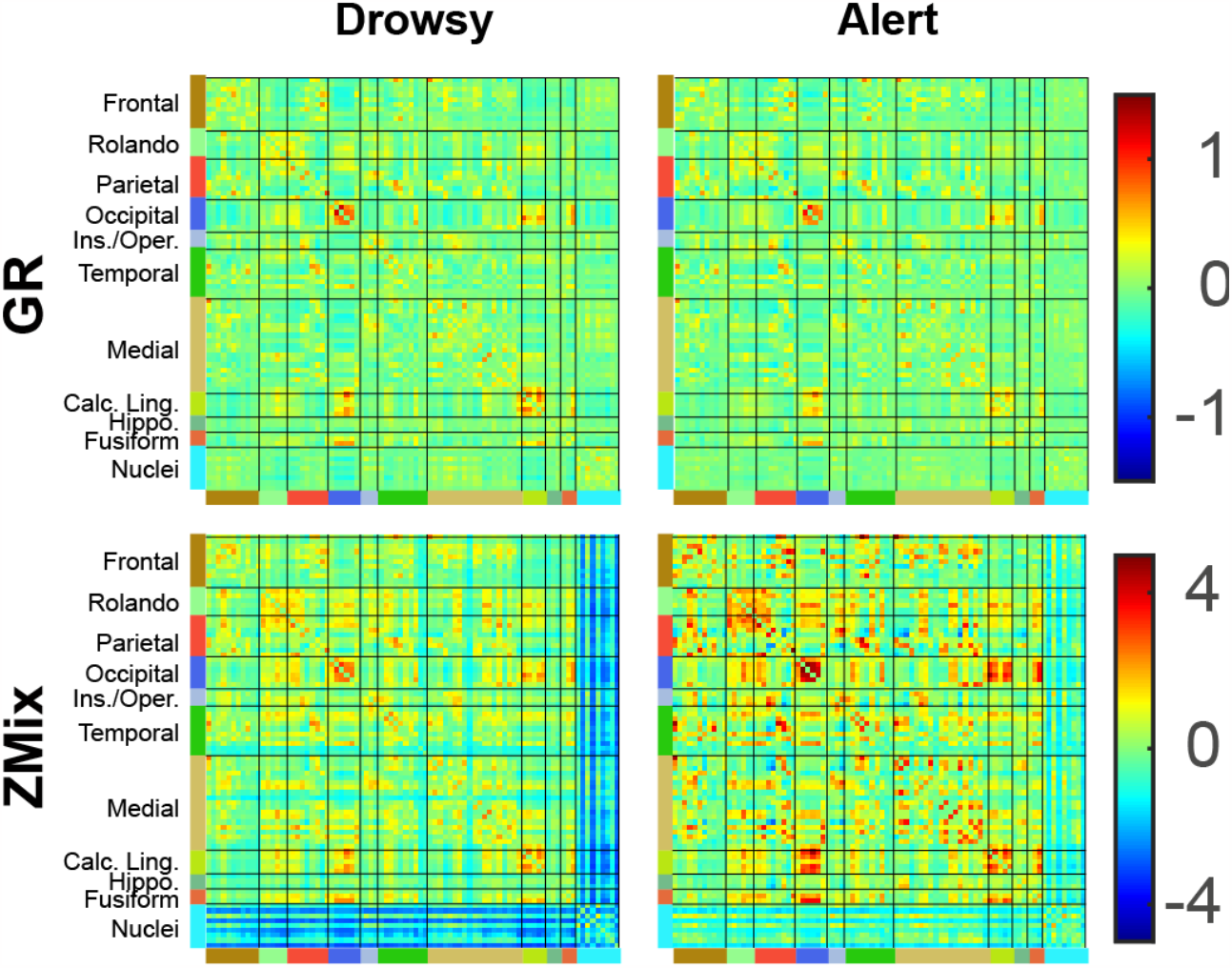
Average z-score intrinsic connectivity computed for drowsy and alert subjects using GR and ZMix corrections. Vigilance Effect on Functional Connectivity

## Discussion

The synthesis analysis of the five methodologies, as shown in Figure 7, provides a comprehensive overview of the differences observed between drowsy and alert subjects across these methodologies. We selected a four-class partition, which roughly corresponds to cortical divisions in the sensory, executive, and Default Mode regions and deep nucleus classes. This partition resembles the gradient-based decomposition proposed by Margulies (2016).

**Fig. 7:**
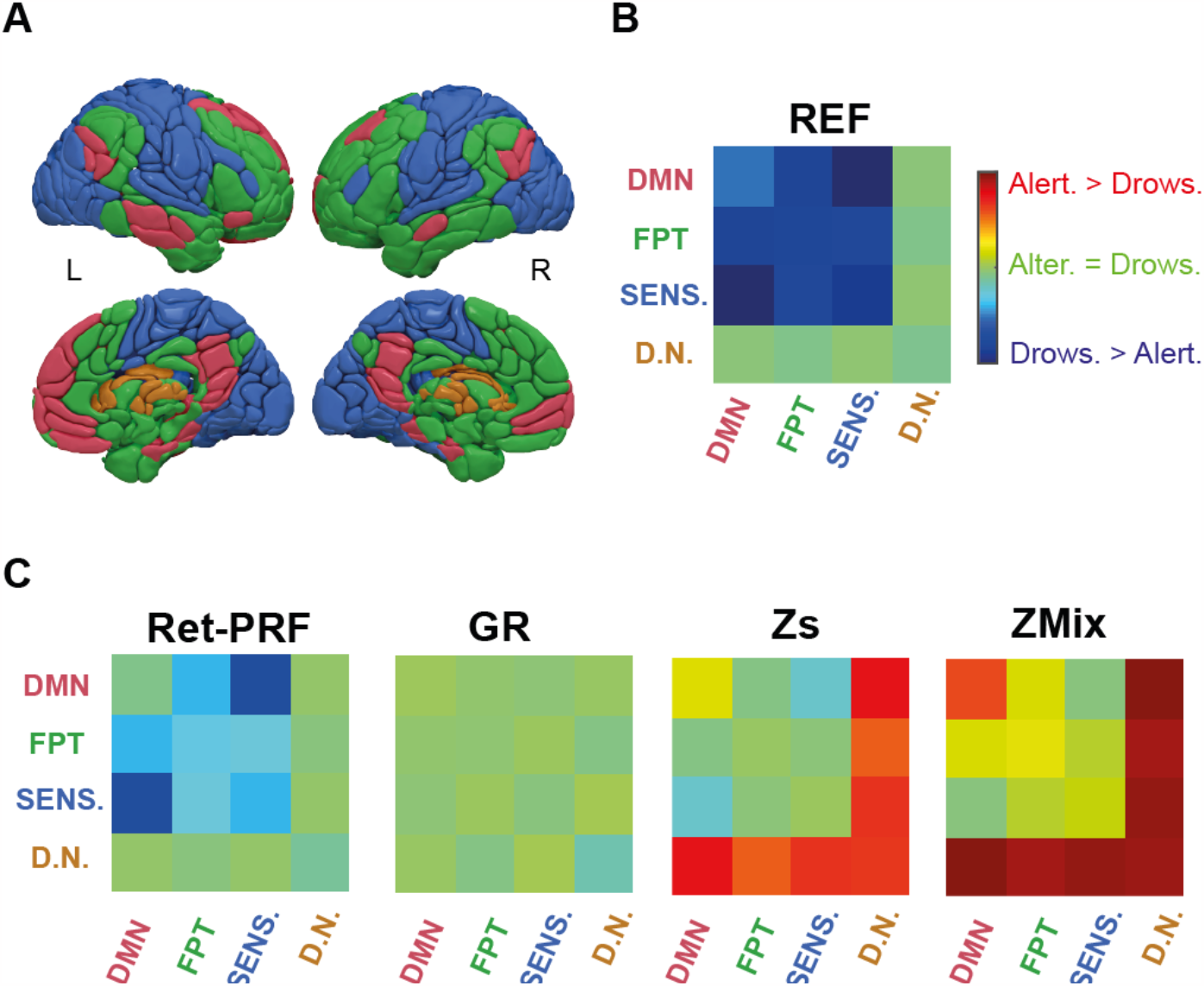
A) Hierarchical decomposition-derived parcellation into four classes (Default mode network/red, Fronto-Parieto-Temporal/green, Sensory related/blue, Deep nuclei/brown). B) Significant vigilance effect on the intra-and inter-class functional connectivity (FCcs) for the reference analysis (REF). C) Significant vigilance effect FCcs for the four types of corrections: Ret-PRF, GR, Sz, and ZMix.

Firstly, our analysis reveals a progression in the functional connectivity patterns from deep nucleus to cortical regions. Initially, the REF analysis showed no significant differences, but as we moved toward the ZMix analysis, there was a shift toward a predominance of alert subjects. Secondly, the cortico-cortical FCs demonstrated a similar evolution. In the REF analysis, drowsy subjects had a more substantial predominance, while in the ZMix analysis, there was a weaker predominance for alert subjects.

Furthermore, the default mode network (DMN) exhibited a distinct pattern that followed the cortico-cortical evolution but displayed a significant predominance in the alert state during the ZMix analysis.

Lastly, our analysis confirms that the GR analysis is the most effective methodology for effectively mitigating the effects of drowsiness.

Various technologies have been developed to mitigate sources of variance associated with movement and susceptibility effects, as summarized in the review by Caballero-Gaudes et al. (Caballero-Gaudes and Reynolds, 2017). These technologies aim to reduce noise and enhance the quality of fMRI data through approaches such as spatial smoothing or region of interest (ROI) analysis. The basic process also involves temporal registration and stereotaxic normalization of fMRI data to a surface or volume atlas. However, temporal slice registration was not applicable in our dataset due to a short repetition time and multiband acquisition, rendering the correction meaningless. Our focus on deep nuclei behavior also led us to choose volumetric stereotaxic normalization. These preprocessing steps led to the “reference” (REF) analysis, from which we deliberately excluded the physiological signal-based corrections that were explicitly studied (see below for discussion).

Our study was motivated by analyzing the resting-state auto-questionnaire completed by all subjects in the MRi-Share cohort (Cremona et al., 2022). This questionnaire included questions directly related to vigilance, such as falling asleep during the acquisition, and indirectly related questions about the number of hours of sleep preceding the acquisition. The responses indicated that 50% of the participants reported some drowsiness, suggesting that vigilance was an issue in this dataset. However, we refrained from using these indexes to classify the two groups, as drowsiness is not always accurately remembered (Ogilvie and Wilkinson, 1984; Hori et al., 1994), and some subjects may choose not to report behaviors that contradict the instructions. The classification of sleep stages is typically based on polysomnography divided into 30-second windows (AASM). Still, this approach may be less suitable for assessing rapid alternations between wakefulness and light sleep observed during drowsiness (Ogilvie and Wilkinson, 1984), especially in the context of an fMRI examination (Tagliazucchi and Laufs, 2014). Instead, we identified drowsiness using a specific vegetative marker: oscillating breathing, which can be observed when a subject fall asleep and disappears quickly as sleep becomes well established (Trinder et al., 1992).

In our study, the mean frequency analysis revealed differences between the drowsy and awake populations in oscillations with a period of approximately 30 seconds, consistent with the characteristics of oscillatory breathing observed at sea level (Khoo et al., 1982). Furthermore, the respiratory rate, heart rate, and estimated respiratory volume per minute were significantly lower in the population exhibiting oscillatory breathing. In contrast, the cardiac variability of this same population was higher. These observations align with the vegetative variations associated with sleep (Benarroch, 2019), confirming that this population experiences a higher level of drowsiness than those without oscillating breathing.

The reference analysis yielded two main findings. First, drowsy subjects generally exhibited higher functional connectivity values than alert subjects. As shown in Figure 3a, this increase varied across drowsy individuals, resulting in an enlarged average histogram for the drowsy group. This variability may reflect different patterns and durations of drowsiness experienced during the 15 minutes of the experiment. Second, no FC wakeful above drowsy functional connectivity group difference was significant, whereas 70% of the drowsy above wakeful differences were significant. These results align with previous findings by Tagliazucchi ((Tagliazucchi and Laufs, 2014)) regarding the difference between sleep stage 1 and wakefulness, as well as the observed differences in functional connectivity between sleepy and alert conditions (Figure 2 of Wang et al., 2017) and wavelet-based analysis of cortico-cortical connections (Spoormaker et al., 2010). They are also consistent with the overall increase in network connectedness observed by Larson-Prior et al. (Larson-Prior et al., 2011). Nguyen et al. (Nguyen et al., 2018), using functional near-infrared imaging, also reported a significant increase in correlation during sleep stage 1 compared to quiet wakefulness, with sleep stage 2 being more similar to wakefulness than sleep stage 1, which contrasts with the findings of Tagliazucchi et al. (Tagliazucchi and Laufs, 2014).

By applying the state-of-the-art physiological-based correction method, Ret-PRF (Glover et al., 2000 Kassinopoulos and Mitsis, 2019), the differences between the two states decreased but remained significant in all cortical regions. This technology corrects both the noise created by the heart and respiration (Glover et al., 2000) and the hemodynamic responses to locally occurring neural processes (Kassinopoulos and Mitsis, 2019). However, this correction method does not capture local spatial specificities accurately, which is a common challenge in task-based fMRI analyses, where the hemodynamic response function is often assumed to be a fixed function (e.g., SPM hemodynamic response function). Nonetheless, the level of correction provided by the respiratory-derived regressor was significantly greater in the drowsy group than in the alert group, confirming that the drowsy state involves neural-based modifications of the respiratory function reflected in the fMRI signal.

Global regression-based correction proved the most effective among the three correction methodologies tested. Only 3.3% of the independent components (FCs) remained significantly different between the two states. Contrary to the controversy, Murthy et al. (Murphy and Fox, 2017) concluded that the use of such correction methods is not inherently “right or wrong” but depends on the scientific question at hand. If the goal is not to study sleep stages, drowsiness, a common confound in resting-state studies (Tagliazucchi and Laufs, 2014), should be accounted for. Global regression-based correction is currently the most effective approach for reducing variability in intrinsic connectivity associated with the drowsy state and making the data of drowsy subjects comparable to data from alert subjects. The other two correction methodologies were designed based on the observation that drowsiness impacts the entire distribution of intrinsic connectivity. Therefore, a global correction approach could be applied. In the case of the pseudo-Gaussian distribution, the most parsimonious correction method is the z-score-based transformation (Sz). This correction equalizes the individual distributions but also shifts the maximum values of the alert subject distribution toward negative values. On the other hand, the mixture model-based correction (ZMix) employs a more complex modeling approach that perfectly aligns the drowsy and alert distributions in both value and amplitude. With the ZMix correction, most differences between the two states manifest as greater FCs in the alert state, with 70% of those including a deep nucleus. For the thalamus (included in 27% of the FCs), several studies have reported a loss of FCs with cortical regions during early NREM sleep (reviewed in Tagliazucchi and van Someren, 2017). The significant predominance of the FCs intra-DMN in the alert state aligns with the expected role of the default mode network compared to other executive or sensorial networks at rest (Raichle et al., 2001, Mazoyer et al., 2001). Moreover, it is consistent with the decoupling observed during NREM sleep (Horovitz et al., 2009; Samann et al., 2011; Larson-Prior et al., 2011; Spoormaker et al., 2012, Wu et al., 2012).

One of the striking messages of this study is that depending on the processing methods used (Ret-PRF, GR, Zs, or Zmix), the interpretation, in terms of networks, of the effects of sleepiness can drastically change or even be reversed without knowing which is the best technique. There is currently no absolute answer, however, the classical data of sleep electrophysiology allow us to propose a framework for reflection.

Studies based on electrophysiological data clearly show that falling asleep is progressively accompanied by a relaxation of the muscles and sensory isolation of the brain to allow the descent into the deeper stages (see for review Ogilvie and Wilkinson, 1984). These functions are under the control of cortico-subcortical loops controlling sensory input via the thalamus and motor control via the basal ganglia. From a connectivity point of view, the progressive isolation of the cortex from the world should decrease intrinsic connectivity between the deep brain nuclei and the cortex in sleepy subjects. This picture is observed with the Zs and Zmix corrections (Figure 7.C).

Moreover, the drowsy subjects observed during an rs-fMRI examination alternate rapidly between a waking state and an N1 stage, contrary to those observed in classical sleep studies where the N1 stage rapidly gives way to deeper stages. This alternation can induce notable hemodynamic variations. Indeed, the descent into sleep is accompanied by vegetative phenomena such as a decrease in heart and respiratory rate or an increase in CO2 partial pressure (Benarroch, 2019). In contrast, micro-arousals, especially during napping, are accompanied by a sympathetic discharge causing tachycardia and peripheral vasoconstriction, among other things (Attoh-Mensah et al., 2023). Thus, subjects who doze off in MRI show cardiorespiratory variations that can alter the MRI signal (Duyn et al., 2020; Claron et al., 2023). Finally, these global and periodic variations in phase with rapid sleep-wake alternations can simulate cortico-cortical pseudo connectivity.

To summarize, the differences in functional connectivity observed between drowsy and alert subjects concerning different networks according to the correction techniques used reflect other mechanisms. On the one hand, the differences in intrinsic connectivity, mainly cortico-cortical, observed between the two populations in the REF condition, which persist after the Ret-PRF correction and which disappear after the GR correction, are primarily a result of the iterative cardiorespiratory variations induced by the vigilance fluctuations. On the other hand, the differences in connectivity concerning cortico-subcortical relationships that appear after the Zs and Zmix corrections reflect the progressive isolation of the cortex from the deep structures observed during the descent into sleep.

## Limitations

The primary limitation of this study stems from the absence of an EEG-based marker for classifying sleep stages. This was due to the challenges of simultaneously acquiring EEG signals during fMRI scans. Incorporating EEG measurements would have prolonged the acquisition time beyond what was deemed acceptable for student participants. However, while the standard classification (Berry et al., 2017) is particularly well suited to the study of sleep, it is less effective at capturing the dynamics of sleep onset, especially if subjects are instructed not to fall asleep ({Ogilvie, 2001 #6200}, Stevner et al., 2019). In addition, oscillatory respiration has proven to be a valuable marker for drowsiness, but it is not guaranteed to be observed in all subjects, particularly in older individuals. Additionally, specific parameters of the scanner’s physiological signal acquisition system, such as the gain, could not be adjusted, resulting in the exclusion of 16% of the subjects. Consequently, due to the lack of sleep stage characterization, the processing methods employed in this study may have underestimated the number of drowsy subjects. There is potential for improvement in this regard, particularly if it becomes feasible to accurately identify « light » sleep periods, which comprise most of the drowsiness period.

## Conclusion

In conclusion, our study demonstrates that the alternation between drowsy and alert phases during the classical resting state paradigm significantly influences the variability of functional connectivity. Among the different processing methodologies, a global regression is a practical approach for reducing this variability. However, the Gaussian-based normalization technique yields the most accurate fit when comparing the functional connectivity between drowsy and alert subjects to the traditional sleep neurophysiological framework.

## Funding

The i-Share cohort has been funded by a grant ANR-10-COHO-05-01 (P.I. C Tzourio) as part of the Programme pour les Investissements d’Avenir. Supplementary funding was received from the Conseil Régional of Nouvelle-Aquitaine, Reference 4370420 (P.I. C Tzourio). The MRi-Share cohort and the ABACI software development have been supported by grants ANR-10-LABX-57 (P.I. B Mazoyer) and ANR-16-LCV2-0006 (GINESISLAB for the software, P.I. M Joliot). The drowsiness study was funded by a grant from France Life Imaging (program ANR-11-INBS-0006, P.I. O. Etard & M. Joliot).

## Notes

### Competing Interest Statement

The authors have declared no competing interest.

